# Chemical convergence between a guild of facultative myrmecophilous caterpillars and host plants

**DOI:** 10.1101/2020.06.29.178319

**Authors:** Luan Dias Lima, José Roberto Trigo, Lucas Augusto Kaminski

## Abstract

Ants exert a strong selective pressure on herbivorous insects, although some caterpillars can live in symbiosis with them using chemical defensive strategies.
We investigated the adaptive resemblance of cuticular hydrocarbons (CHCs) in multitrophic systems involving a guild of facultative myrmecophilous caterpillar species (Lepidoptera: Lycaenidae), tending ants (Hymenoptera: Formicidae) and host plants from three families. We hypothesized that the CHCs of the caterpillars would resemble those of their host plants (chemical camouflage).
We analyzed CHCs using gas chromatography/mass spectrometry. Morisita’s similarity index (SI) was used to compare CHC profiles of caterpillar species with different types of ant associations (commensal or mutualistic), ants and host plants.
We found strong convergence between caterpillars’ CHCs and plants, especially for commensal species that do not provide secretion rewards for ants. Moreover, we found unexpected chemical convergence among mutualistic caterpillar species that offer nectar reward secretions to ants.
These results show that the studied caterpillars acquire CHCs through their diet and that they vary according to host plant species and type of ant association (commensalism or mutualism). This ‘chemical camouflage’ of myrmecophilous caterpillars may have arisen as a defensive strategy allowing coexistence with ants on plants, whereas ‘chemical conspicuousness’ may have evolved in the context of honest signaling between true mutualistic partners.
We suggest the existence of both Müllerian and Batesian chemical mimicry rings among myrmecophilous caterpillar species. Cuticular chemical mixtures can play a key adaptive role in decreasing ant attacks and increasing caterpillar survival in multimodal systems.

**Graphical abstract:**

- Chemical camouflage can be a defensive strategy of myrmecophilous caterpillars against ants.
- ‘Chemical conspicuousness’ is proposed as a new strategy mediated by cuticular hydrocarbons in myrmecophilous caterpillars.
- Chemical mimicry rings can occur between myrmecophilous caterpillars and especially between mutualistic species that produce nectar rewards for ants.

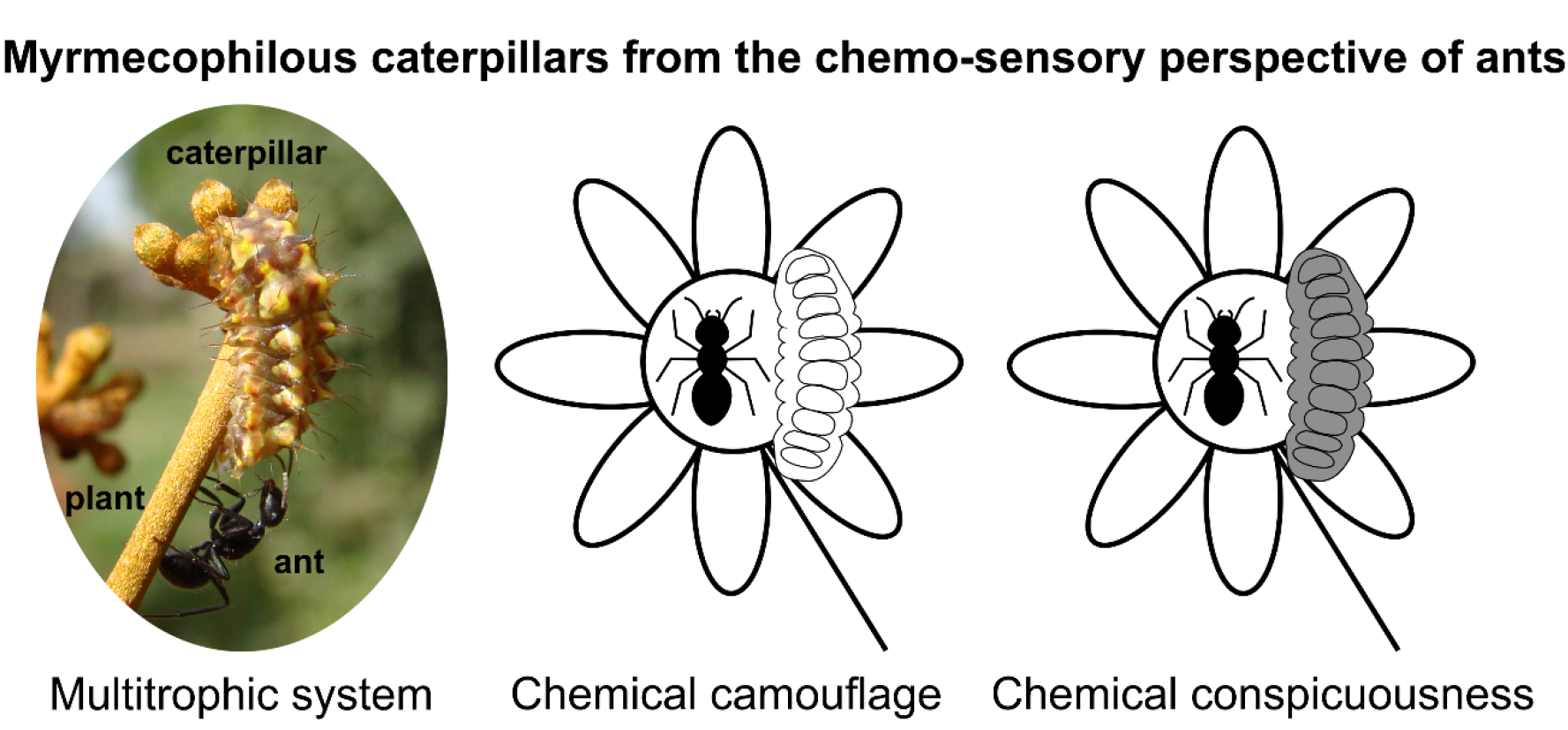

## Introduction

Herbivorous insects suffer a top-down effect from predators and parasitoids (Vidal & Murphy, 2018). Strategies that minimize detection or attack after detection, as well as deceive these natural enemies, will be positively selected and spread within herbivorous insect species (see Ruxton *et al.*, 2004). Among top predators in terrestrial ecosystems, ants patrolling foliage are considered to apply strong selective pressure on herbivorous insects (Floren *et al.*, 2002). However, some myrmecophilous herbivores, such as hemipterans (e.g. aphids, scales, coccids, whiteflies, leafhoppers, and treehoppers), lepidopteran caterpillars (Lycaenidae and Riodinidae), and aphids and cynipid wasps (via galls), can release liquid rewards rich in sugar (honeydew) that attract tending ants (see Pierce *et al.*, 2002; Pierce, 2019; Pringle, 2020). Such a symbiotic relationship between ants and the trophobiont insects that they attend is termed trophobiosis (Gibernau & Dejean, 2001). The attraction of tending ants can protect trophobiont herbivores against predators and parasitoids as a conditional mutualism since ants patrolling foliage behave aggressively against any foreign arthropod (see Rico-Gray & Oliveira, 2007 and references therein for examples with hemipterans).

Trophobiosis is particularly interesting in lycaenid and riodinid caterpillars. These caterpillars do not excrete the excess of sugar-rich phloem as honeydew-producing hemipterans do, but instead produce costly secretion rewards through specialized glands (Daniels *et al.*, 2005; Kaminski & Rodrigues, 2011). This secretion of lycaenid caterpillars has been reported to manipulate the behavior of tending ants (Hojo *et al.*, 2015). Furthermore, myrmecophilous caterpillars possess behavioral traits and sets of specialized ‘ant-organs’ used for chemical and acoustic communication with tending ants (Malicky, 1970; DeVries, 1990; Fiedler *et al.*, 1996; Casacci *et al.*, 2019). However, a question remains: Is the nutritional reward offered by trophobiont caterpillars sufficient to deter ant aggressive behavior, or have additional defensive mechanisms evolved to avoid predation? Several studies have shown that tending ants may prey upon honeydew-producing aphids (e.g. Sakata *et al.*, 1995; Fischer *et al.*, 2001). Thus, an alternative strategy may be ‘chemical camouflage’ (sensu von Beeren *et al.*, 2012), by which an operator does not detect an emitter because its chemical cues blend with the environment and, thus, no reaction is caused in the operator. This strategy is found in the myrmecophilous treehopper *Guayaquila xiphias* (Fabricius) (Hemiptera: Membracidae) due to the high degree of similarity between its cuticular hydrocarbons (CHCs) and its host plant *Schefflera vinosa* (Cham. & Schltdl.) Frodin & Fiaschi (Araliaceae) and seems to solve the dilemma of attracting aggressive tending ants, since these predators fail to recognize the hemipteran as prey (Silveira *et al.*, 2010). Thus, we aimed to investigate the adaptive resemblance among CHC profiles in three multitrophic systems involving a guild of facultative myrmecophilous caterpillars, tending ants and host plant species (Fig. 1). These caterpillars are polyphagous with color pattern associated with visual camouflage (polychromatism) (see Monteiro, 1991). Thus, we hypothesized that these caterpillars would have CHC profiles that resemble those of their host plants, suggesting chemical camouflage as an additional defensive mechanism against aggressive ants.

**Fig. 1.**
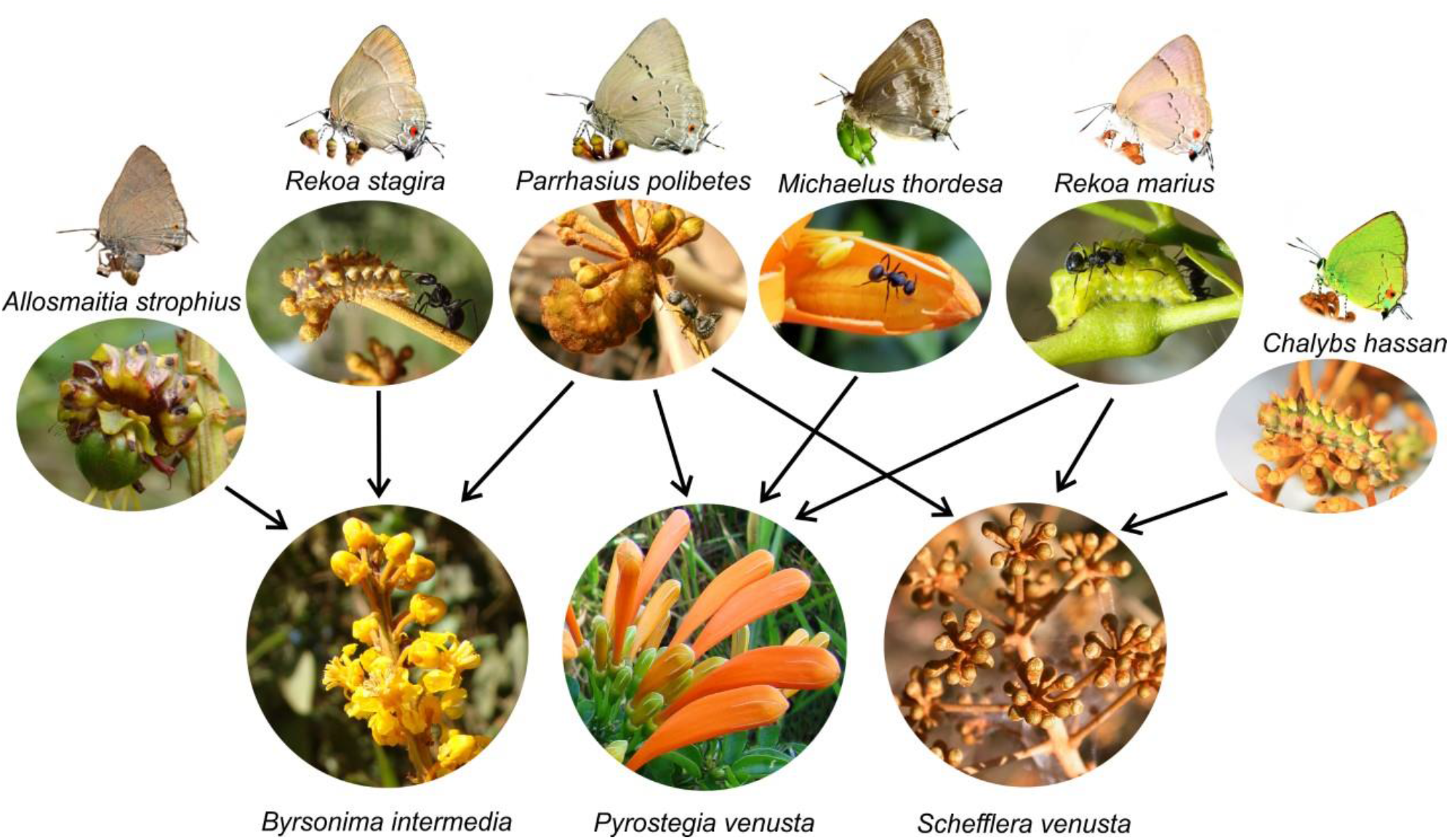
Multitrophic systems studied involving a guild of six facultative myrmecophilous caterpillars (Lepidoptera: Lycaenidae), tending ants (Hymenoptera: Formicidae) and host plant species of three families (Malpighiaceae, Bignoniaceae and Araliaceae) respectively. Photos of *Michaelus thordesa* by Sabrina Thiele ©

## Materials and methods

### Study Site and Organisms

Collections were carried out in two areas of Cerrado (Brazilian savanna) in the state of São Paulo, in Southeast Brazil: a small fragment belonging to Laboratório Nacional de Luz Síncrotron in the city of Campinas (22°48’S, 47°03’W); and a reserve of Estação Experimental de Mogi-Guaçu in the city of Mogi Guaçu (22°18’S, 47°10’W). The vegetation of both sites consists of cerrado *sensu stricto*, a dense scrubland of shrubs and trees (Oliveira-Filho & Ratter, 2002). The collection and transport of the specimens were authorized by the Sistema de Autorização e Informação em Biodiversidade (SISBIO) by the license No. 62345-1.

Neotropical lycaenid butterflies are included in three subfamilies, within which Eumaeini is the most diverse tribe (Robbins, 2004). Eumaeini caterpillars are generally polyphagous and feed on reproductive tissue (buds and flowers) of host plants (see Robbins & Aiello, 1982; Kaminski *et al.*, 2012; Silva *et al.*, 2014). These caterpillars are engaged in low-degree facultative interactions with tending ants, although several species seem to have lost myrmecophily (Fiedler, 1991; LAK, unpublished data). Immatures (eggs and caterpillars) of eumaeine species were collected from three host plant species that are commonly used by these caterpillars at the study sites (see Kaminski & Freitas, 2010; Kaminski *et al.*, 2010b, 2012; Rodrigues *et al.*, 2010): *Byrsonima intermedia* A. Juss. (Malpighiaceae), *Pyrostegia venusta* (Ker-Gawl.) Miers (Bignoniaceae) and *Schefflera vinosa* (Araliaceae). Six eumaeine species were collected from the inflorescences of these plants (Fig. 1): (1) *Allosmaitia strophius* (Godart) is an oligophagous caterpillar specialized on inflorescences of Malpighiaceae (Kaminski & Freitas, 2010). Although females prefer to oviposit on plants with ants (Bächtold *et al.*, 2014), the caterpillars have a non-functional dorsal nectary organ (DNO) — that is, they do not establish a typical food-for defense mutualistic interaction with ants, and thus are considered commensal myrmecophilous (i.e. an organism that indirectly interacts with ants) (Kaminski & Freitas, 2010; Silva *et al.*, 2014). (2) *Chalybs hassan* (Stoll) is a polyphagous caterpillar on flower buds of Araliaceae, Fabaceae, Sapindaceae, and Sterculiaceae (LAK, unpublished data). They possess a DNO, but there is no evidence regarding its functionality and role in myrmecophily. (3) *Michaelus thordesa* (Hewitson) is a polyphagous caterpillar, but occurs most frequently on tubular flower buds of Bignoniaceae, and lives inside of flowers. It is a facultative myrmecophilous with a functional DNO (Silva *et al.*, 2014; Thiele & Kaminski, in prep.). (4) *Parrhasius polibetes* (Stoll) is a polyphagous caterpillar that is facultative myrmecophilous with a functional DNO beginning in the third instar, while it is commensal myrmecophilous in the first and second (Kaminski *et al.*, 2010a, 2012; Rodrigues *et al.*, 2010). (5) *Rekoa marius* (Lucas) is a polyphagous caterpillar that is facultative myrmecophilous with functional DNOs on the third and last instars (Monteiro, 1991; Silva *et al.*, 2014; Faynel *et al.*, 2017). (6) *Rekoa stagira* (Hewitson) is a polyphagous caterpillar that is facultative myrmecophilous with functional DNOs on the third and last instars (Faynel *et al.*, 2017; LAK, unpublished data).

Samples of two ant species, *Camponotus blandus* (Smith) and *Camponotus crassus* Mayr were collected from the host plants. These species are very common in the Cerrado and are involved in most of the interactions between ants and trophobionts on plants, including myrmecophilous caterpillars (e.g. Alves-Silva *et al.*, 2013; Bächtold, 2014; Lange *et al.*, 2019).

The hatched eggs and caterpillars were reared on buds or flowers of their respective host plant in an uncontrolled environment in the laboratory. For chemical analysis, the desired developmental stage (different larval instars) of the butterflies, the buds or flowers of host plants, and the tending ants were killed by freezing and kept frozen at −20°C until the CHC extraction. When available, both the second instar, which does not produce a reward secretion, and the fourth instar (last instar from now on), which may produce one, were collected. The following samples were collected for analysis: *B. intermedia* (N = 2 bud, N = 1 flower); *P. venusta* (N = 1 bud, N = 1 flower); *S. vinosa* (N = 1 bud); *A. strophius* (N = 2 last instar on *B. intermedia*); *C. hassan* (N = 1 last instar on *S. vinosa*); *M. thordesa* (N = 1 last instar on *P. venusta*); *P. polibetes* (N = 1 second instar, N = 2 last instar on *B. intermedia*, N = 4 last instar on *P. venusta,* and N = 1 last instar on *S. vinosa*); *R. marius* (N = 1 last instar on *P. venusta* and N = 1 last instar on *S. vinosa*); *R. stagira* (N = 1 last instar on *B. intermedia*); *C. blandus* (N = 1 pool of 20 workers) and *C. crassus* (N = 1 pool of 20 workers).

### Extraction and Identification of Cuticular Compounds

CHCs of the organisms were extracted following Portugal & Trigo (2005). The organisms were dipped in 5 ml of *n*-hexane (95%, Ultra-Resi-Analysed J.T.Baker) for 5 min and then removed with a forceps. The hexane was subsequently treated with anhydrous Na_2_SO_4_, filtered, and evaporated gently in a stream of N_2_. The CHC extracts were analyzed using electron impact gas chromatography–mass spectrometry in a gas chromatograph (Hewlett Packard 6890) equipped with HP-5MS column (5% phenyl methyl siloxane capillary 95%, 30 m × 250 μm × 0.25 μm; Hewlett Packard) directly coupled to a mass selective detector (Hewlett Packard 5973). All analyses were performed under the following conditions: 250°C temperature of injection; 60° to 300°C at 2°C/min, 20 min at 300°C program temperature; helium 1 mL/min as carrier gas; ionization energy of 70 eV and a range of 40–600 amu; splitless injection mode, 1 μl injected. All samples were analyzed with and without co-injection with consecutive *n*-alkane standards for the determination of Kovats Retention Index (KI). Alkanes and alkenes were identified by their KI (Carlson *et al.*, 1998) and fragmentation patterns (Nelson *et al.*, 1981; Pomonis, 1989; Carlson *et al.*, 1999; Howard, 2001). Alkene identification was confirmed after derivatization with dimethyl disulfide for determining the double-bond position (Francis & Veland, 1981). The fragmentation pattern of *n*-alcohols was compared with those in the literature (Wang *et al.*, 2007). The identification of *n*-alcohols was confirmed after derivatization with *N*-methyl-*N*-trimethylsilytrifluoruoacetamide (MSTFA; 100 mL MSTFA, 80°C, 1 h) according to Menéndez *et al.* (2005). Some cuticular compounds were tentatively assigned (e.g. squalene-like) using the NIST Mass Spectral Search Program (Agilent Technologies, Version 2.0 f. 2008) together with mass fragmentation interpretation of Budzikiewicz *et al.* (1967). The remaining compounds remained as unknowns.

### Statistical Analyses

To test our hypothesis, we calculated percentages of absolute abundance of the compounds found in the cuticular extracts by considering the compounds as 100%. These data were then used to calculate relative abundances, that is, the quantity of each separate compound expressed as a percentage of the total occurrence of the class of substance. We performed a cluster analysis using Morisita’s similarity index (SI) to compare CHC profiles of the studied species. This index varies from 0% (no similarity) to 100% (complete similarity) (Krebs, 1999). We used an analysis of similarity (ANOSIM) to test significant differences based on the percentage of similarity of the CHC profiles between species. Data were partitioned into two separate analysis to facilitate comparisons. First, the species were divided into three groups: caterpillars (group 1), ants (group 2), and plants (group 3). Second, the species were divided into four groups: commensal caterpillars (group 1), mutualistic caterpillars (group 2), ants (group 3), and plants (group 4). In these analyses, R values from close or equal to 0 (total similarity) to 1 (total difference) were calculated between the groups (see Clarke, 1993). As *C. hassan* did not have any kind of ant association (commensal or mutualistic) confirmed, it was not analyzed in the second comparison. All analyses were performed with PAST software (Version 4.03). Values of similarity above 80% between caterpillars and host plants were considered possible cases of chemical camouflage strategy. This value was defined based on the bioassays carried out by Silveira *et al.* (2010), who showed this value to be sufficient to significantly reduce detection of myrmecophilous insects chemically camouflaged against ants of the genus *Camponotus*.

## Results

Gas chromatography–mass spectrometry analyses of cuticular extracts revealed a high degree of similarity between eumaeine caterpillars and their host plants (R_ANOSIM_ = 0.20; P < 0.05) (Figs. 2–3 and 5; Tables 1 and S1–S5). In general, the caterpillars and their host plants had *n*-alkanes (C27 and C29) as their main components (Figs. 2–3; Tables S1–S3). The highest SI values were between caterpillars that fed on *B. intermedia*, which ranged from 35% to 98%, followed by those that fed on *P. venusta*, ranging from 55% to 90%. Similarity was low for caterpillars on *S. vinosa*, ranging from 23% to 34% (Figs. 2–3 and 5; Table 1). Evidence of chemical camouflage (similarity > 80%) was found for four caterpillar species: *A. strophius* (98%), *P. polibetes* (second instar) (96%) and *R. stagira* (96%) on *B. intermedia*, and *R. marius* on *P. venusta* (90%) (Fig. 5). The highest values of convergence with the host plant were for commensal myrmecophilous that do not have a functional DNO that produces nectar rewards (R_ANOSIM_ = −0.08; P > 0.05), namely *A. strophius* (98%) and the second instar of *P. polibetes* (96%) (Figs. 2–3 and 5; Tables 1 and S5).

**Table 1.** Morisita’s similarity index (mean ± standard error) of the shared cuticular hydrocarbons of lycaenid caterpillars, host plants, and tending ants. Similarity values > 0.8 in bold

| Lycaenid caterpillars | Host plants |  |  |  | Lycaenid caterpillars |  |  |  |  | Tending ants |  |  |
| --- | --- | --- | --- | --- | --- | --- | --- | --- | --- | --- | --- | --- |
|  | *Byrsonima intermedia* | *Pyrostegia venusta* | *Schefflera vinosa* | *Allosmaitia strophius* | *Chalybs hassan* | *Michaelus thordesa* | *Parrhasius polibetes* (S) | *Parrhasius polibetes* (L) | *Rekoa marius* | *Rekoa stagira* | *Camponotus blandus* | *Camponotus crassus* |
| *Allosmaitia strophius* (L)* | **0.93 ± 0.02** | – | – | – | 0.39 ± 0.00 | 0.25 ± 0.01 | **0.85 ± 0.00** | 0.34 ± 0.01 | 0.39 ± 0.05 | **0.95 ± 0.00** | 0.03 ± 0.00 | 0.08 ± 0.00 |
| *Chalybs hassan* (L)** | – | – | 0.34 | 0.39 ± 0.00 | – | 0.76 | 0.54 | 0.78 ± 0.02 | 0.62 ± 0.03 | 0.54 | 0.09 | 0.09 |
| *Michaelus thordesa* (L) | – | 0.55 ± 0.00 | – | 0.25 ± 0.01 | 0.76 | – | 0.36 | **0.80 ± 0.06** | 0.58 ± 0.01 | 0.41 | 0.05 | 0.04 |
| *Parrhasius polibetes* (S)* | **0.90 ± 0.03** | – | – | **0.85 ± 0.00** | 0.54 | 0.36 | – | 0.51 ± 0.03 | 0.54 ± 0.11 | **0.92** | 0.06 | 0.13 |
| *Parrhasius polibetes* (L) | 0.44 ± 0.04 | 0.60 ± 0.02 | 0.27 | 0.34 ± 0.01 | 0.78 ± 0.02 | **0.80 ± 0.06** | 0.51 ± 0.03 | 0.77 ± 0.05 | 0.66 ± 0.01 | 0.51 ± 0.03 | 0.14 ± 0.02 | 0.10 ± 0.01 |
| *Rekoa marius* (L) | - | **0.82 ± 0.08** | 0.23 | 0.39 ± 0.05 | 0.62 ± 0.03 | 0.58 ± 0.01 | 0.54 ± 0.11 | 0.66 ± 0.01 | - | 0.53 ± 0.09 | 0.06 ± 0.01 | 0.15 ± 0.04 |
| *Rekoa stagira* (L) | **0.94 ± 0.02** | - | - | **0.95 ± 0.00** | 0.54 | 0.41 | **0.92** | 0.51 ± 0.03 | 0.53 ± 0.09 | – | 0.03 | 0.10 |

\*Commensal myrmecophily, dorsal nectary organ (DNO) non-functional; L = last instar; S = second instar. \*\*Lack of data on DNO functionality and kind of ant association.

**Fig. 2.**
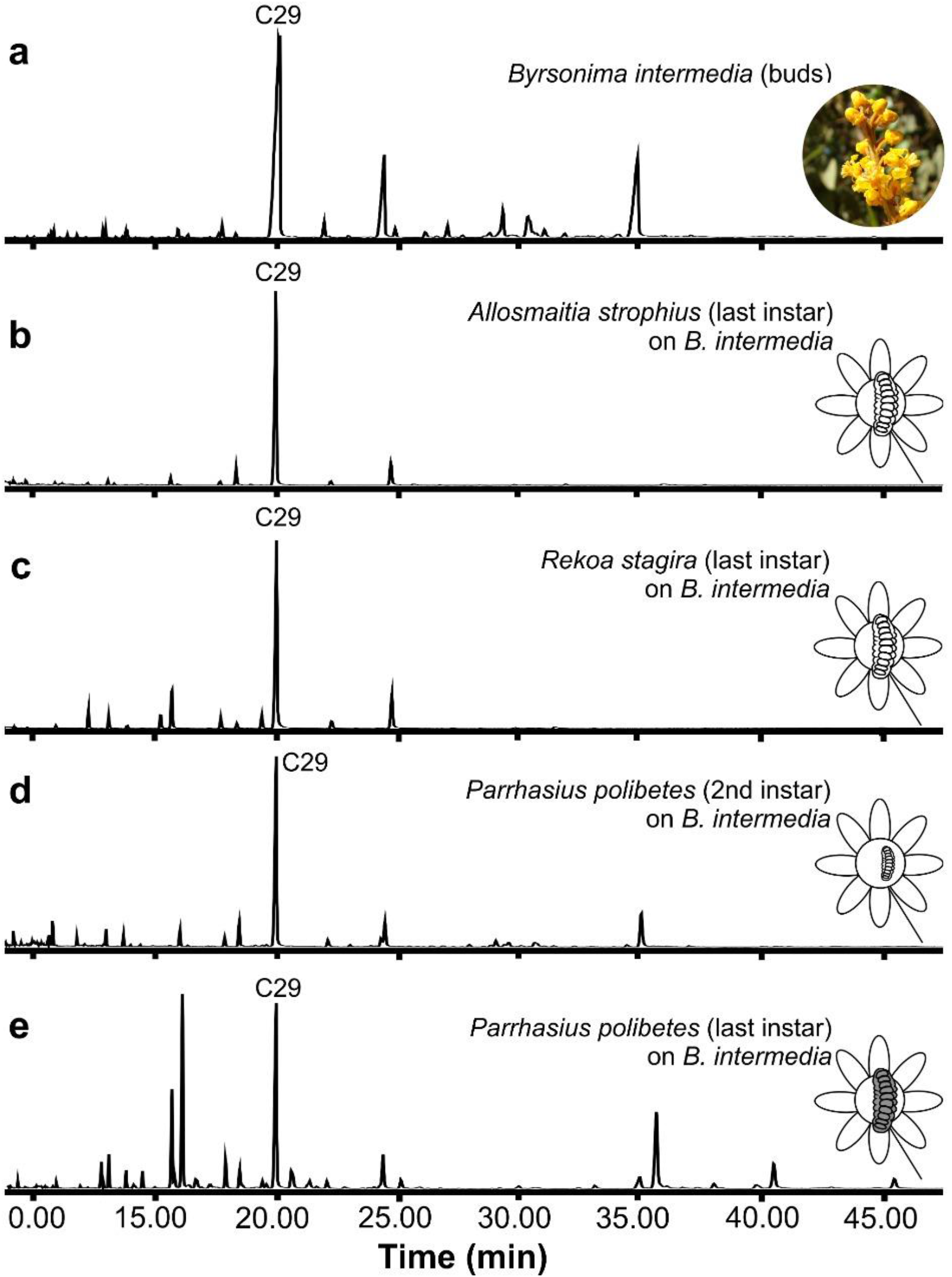
Chromatograms of cuticular compounds of *Byrsonima intermedia* (Malpighiaceae) buds (a) and of associated facultative myrmecophilous caterpillars (Lepidoptera: Lycaenidae) (b-e). (b) last instar of *Allosmaitia strophius*, (c) last instar of *Rekoa stagira*, (d) second instar of *Parrhasius polibetes*, and (e) last instar of *P. polibetes*. Schematic white-colored caterpillars represent cases of the ‘chemical camouflage’ strategy (similarity > 80%) while gray-colored caterpillars represent cases of the ‘chemical conspicuousness’ strategy

Last instars of the mutualistic myrmecophilous *P. polibetes* and *R. marius* have conspicuous and less variable profiles that are less affected by host plant composition. Surprisingly, there were also high SI values between some caterpillar species regardless of the host plant. For example, similarities between the last instars of *M. thordesa* and *P. polibetes* (from 84% to 95%) and last instars of *C. hassan* and *M. thordesa* (76%) were higher than the similarities with their host plants (Figs. 2–3; Table 1). In general, the SI values for the last instar of *P. polibetes* were higher with other caterpillars (25% to 95%) than of the host plant used (27% to 66%) (Figs. 2–3; Table 1). Moreover, the mutualistic caterpillars did not have high similarities with plants (R_ANOSIM_ = 0.55; P < 0.05; Table S5).

In contrast, the chemical similarity between the caterpillars and ants of the genus *Camponotus* was lower and with a different pattern of cuticular compounds (SIs < 23%, ANOSIM R_ANOSIM_ = 1; P < 0.05) (Figs. 4–5; Tables 1, S1, and S4–S5). Worker ants of *C. blandus* and *C. crassus* had mainly branched alkanes and unidentified compounds, with a SI between them of 28% (Figs. 4–5).

**Fig. 3.**
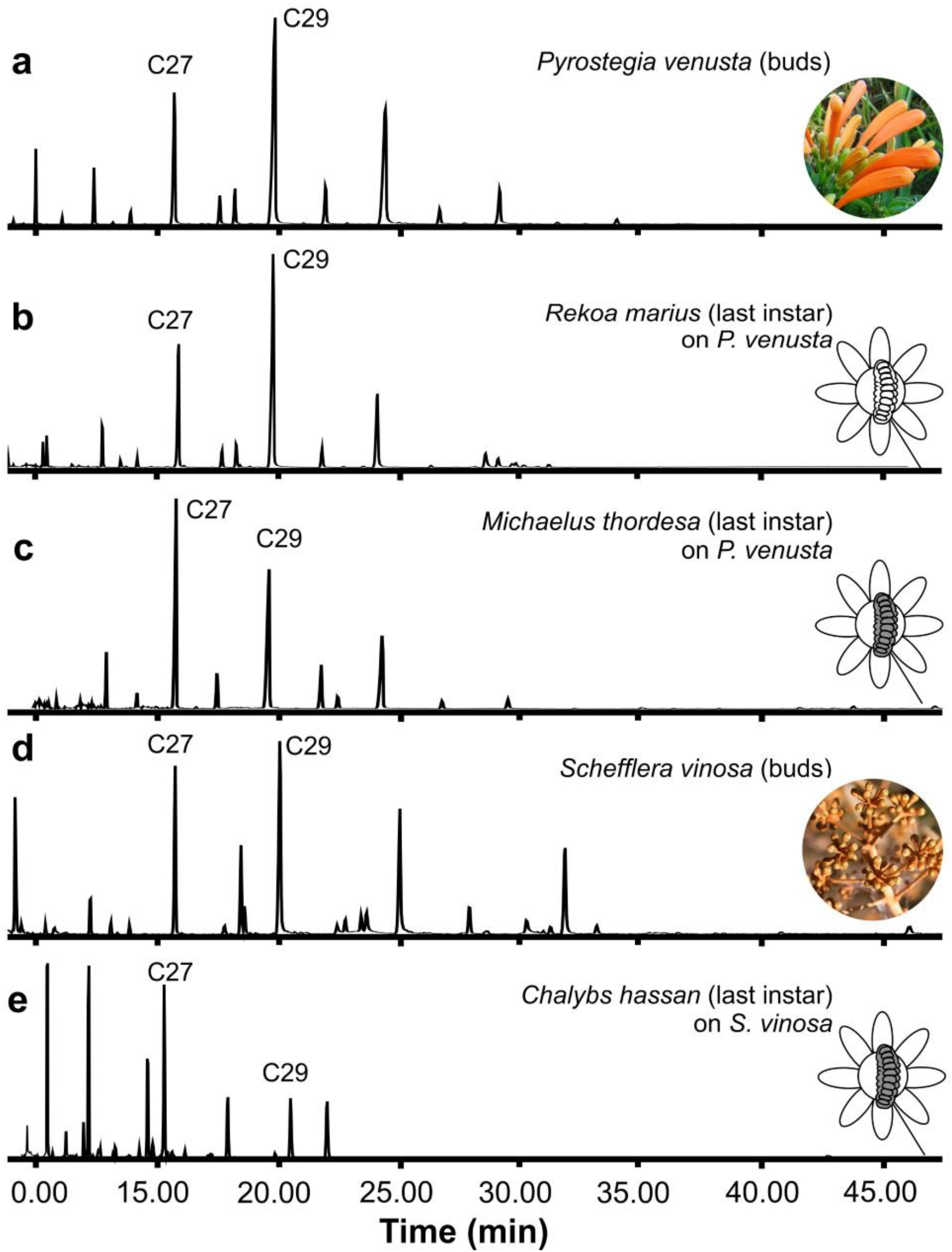
Chromatograms of the cuticular compounds of host plants (a, d) and associated facultative myrmecophilous caterpillars (Lepidoptera: Lycaenidae) (b-c, e). (a) *Pyrostegia venusta* (Bignoniaceae) buds, (b) last instar of *Rekoa marius* on *P. venusta*, (c) last instar of *Michaelus thordesa* on *P. venusta*, (d) buds of *Schefflera vinosa* (Araliaceae), and (e) last instar of *Chalybs hassan* on *S. vinosa*. Schematic white-colored caterpillar represent cases of the ‘chemical camouflage’ strategy (similarity > 80%) while gray-colored caterpillars represent cases of the ‘chemical conspicuousness’ strategy

**Fig. 4.**
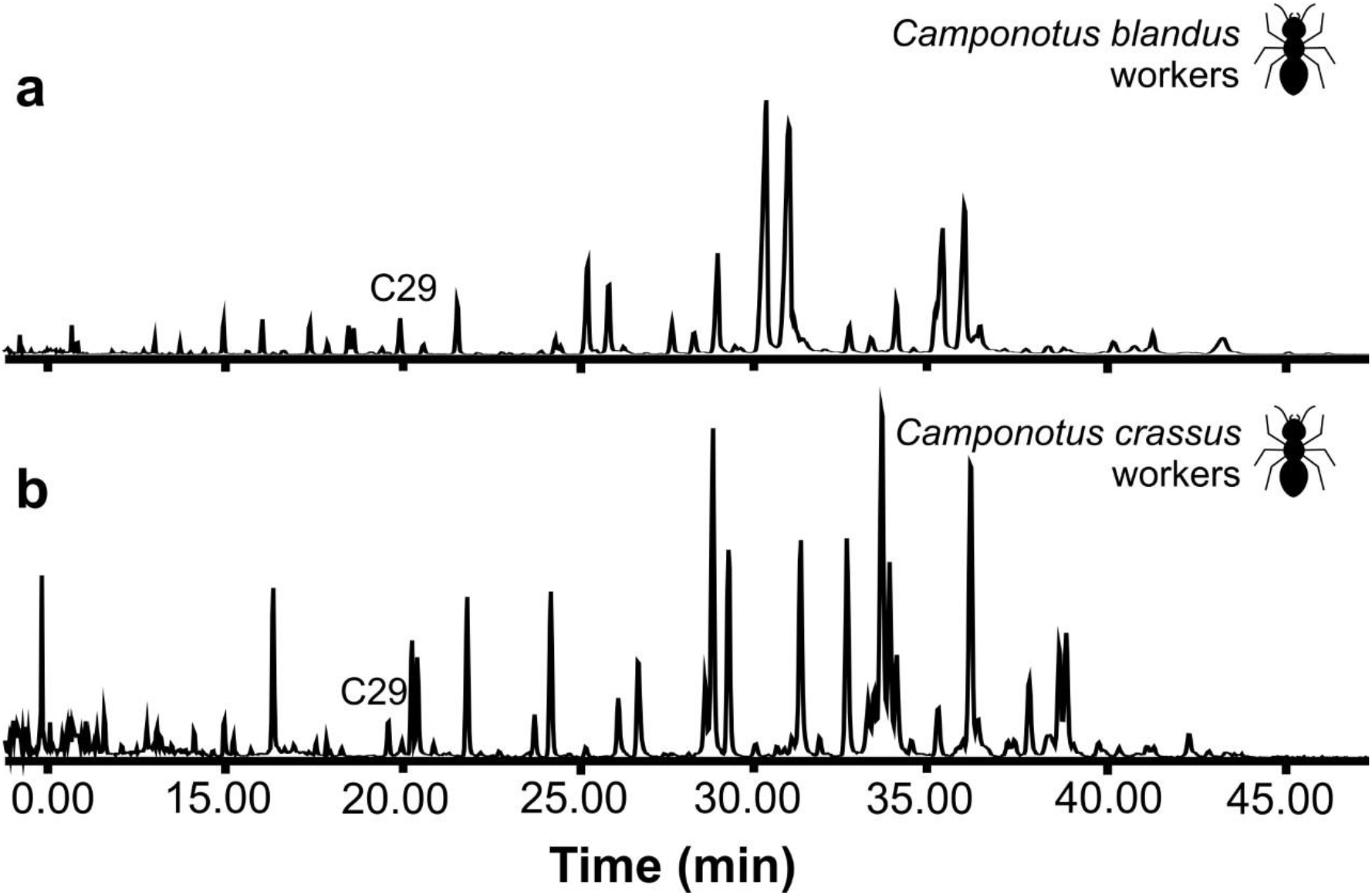
Chromatograms of the cuticular compounds of tending ants (Hymenoptera: Formicidae). (a) *Camponotus blandus* workers, (b) *Camponotus crassus* workers

**Fig. 5.**
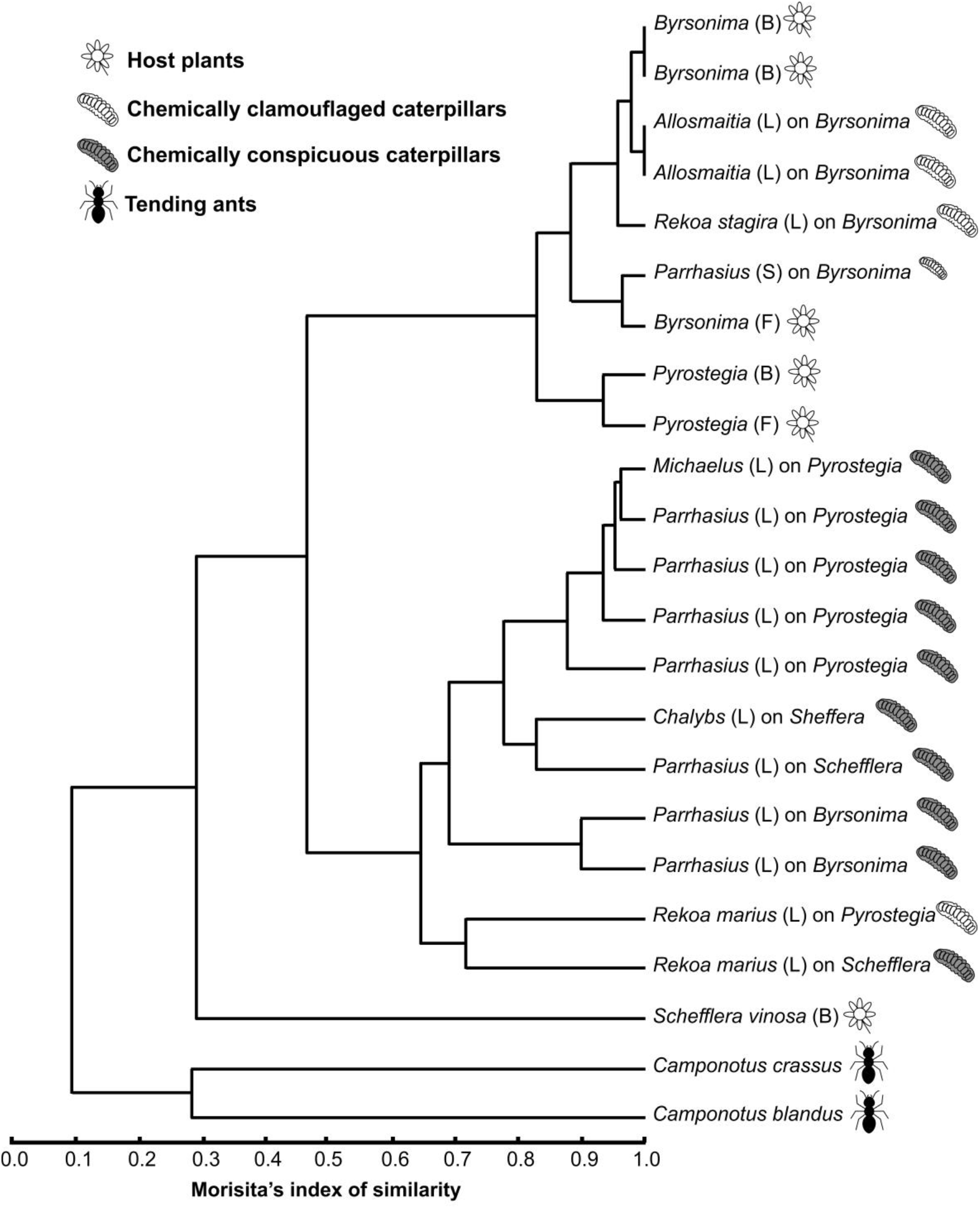
Hierarchical cluster analysis (Morisita’s similarity index) of the shared cuticular hydrocarbons of facultative myrmecophilous caterpillars (Lepidoptera: Lycaenidae), host plants and tending ants. B = bud; F = flower; L = last instar; S = second instar

## Discussion

We found a high degree of similarity between the CHC profiles (> 80%) of some species of myrmecophilous caterpillars and of host plants from two of the plant families analyzed in this study, thus confirming our initial hypothesis. The fact that the CHC profiles of caterpillars were congruent with the profiles of the plants they were feeding on suggests that these caterpillars can acquire these profiles through their diet. It has been shown that the diet can be essential on the chemical camouflage strategy (e.g. Akino *et al.*, 2004; Lohman *et al.*, 2006; Lima *et al.*, in prep.). This kind of chemical camouflage, also referred to as chemical crypsis or phytomimesis, was suggested by Espelie *et al.* (1991) who found similarity between the cuticular lipids of herbivorous insects and their host plants (see Akino *et al.*, 2004; Akino, 2005, 2008; von Beeren *et al.*, 2012; Lima & Kaminski, 2019). The phenomenon was later reported for the first time for a non-trophobiont, namely the caterpillar of *Biston robustum* Butler (Geometridae) (Akino *et al.*, 2004; Akino, 2005). Caterpillars of the butterfly *Mechanitis polymnia* (Linnaeus) (Ithomiinae) and the moth *Cydia pomonella* (Linnaeus) (Tortricidae), as well as larvae of the beetle *Chelymorpha reimoseri* Spaeth (Chrysomelidae) and wasps, also possess this defensive strategy against chemically oriented predators, including ants, for example (Portugal & Trigo, 2005; Piskorski *et al.*, 2010; Massuda & Trigo, 2014; Ranganathan *et al.*, 2015).

This strategy has been shown to prevent ants from recognizing trophobiont treehoppers as prey (Silveira *et al.*, 2010) and to reduce ant attacks even in the absence of honeydew rewards (Wang *et al.*, 2018). However, to our knowledge, within Lycaenidae, chemical camouflage has only been demonstrated for the entomophagous caterpillars of *Feniseca tarquinius* (Fabricius), which have a similar lipid composition to that of their aphid prey, which are mutualistic with ants (Youngsteadt & DeVries, 2005; Lohman *et al.*, 2006). Silveira *et al.* (2010) suggested that chemical camouflage could also occur with ant-tended caterpillars and that this strategy could function as insect analogs of extrafloral nectaries for ants. Thus, this is the first evidence of chemical camouflage among trophobiont caterpillars. It is known that ants may prey on trophobiont species that produce less honeydew (Sakata *et al.*, 1995), and that lycaenid caterpillars can use their secretions to appease the aggressive behavior of ants (Hojo *et al.*, 2015).

Nonetheless, our results suggest that trophobiotic caterpillars can use an additional CHC-mediated strategy through chemical background matching even when they are unable to secrete honeydew. This occurs either because they are in the second instar when their DNOs, which produce nutritious secretions for ants, are not functional (see Fiedler, 1991; Kaminski *et al.*, 2010a) or when they are in the pupal stage when DNOs are not retained (Mizuno *et al.*, 2018). Indeed, some of the highest similarity values between caterpillars and host plants found in the present study were for *A. strophius*, a commensal myrmecophilous that does not produce nectar rewards (Kaminski & Freitas, 2010; Bächtold *et al.*, 2014). The cuticular compounds of the facultative myrmecophilous lycaenid pupa *Lycaeides argyrognomon* (Berstrasser) contains not only CHCs but also several long-chain aliphatic aldehydes that suppress ant aggression even after certain ant-organs are non-functional (Mizuno *et al.*, 2018).

Moreover, myrmecophilous lycaenid caterpillars have several adaptations, such as chemical defense using exocrine glands, hairiness, thickened larval cuticle, and/or construction of silken shelters, that avoid ant aggression (Malicky, 1970; Pierce *et al.*, 2002). They also have ant-associated organs, such as pore cupola organs (PCOs), which supposedly secrete substances to pacify ants, tentacle organs (TOs) and DNO (Pierce *et al.*, 2002). Lycaenid caterpillars with a specialized parasitic lifestyle may use chemical or acoustic mimicry as well (e.g. Nash *et al.*, 2008; Barbero *et al.*, 2009). However, the low degree of similarity between the CHC profiles of the caterpillars and ants studied here discards the hypothesis of this kind of chemical mimicry in facultative myrmecophilous interactions.

On the other hand, lycaenids are also benefited by ant protection and grooming (Hölldobler & Wilson, 1990; Fiedler *et al.*, 1996; Hojo *et al.*, 2014b) and ants may use the non-congruent CHCs of *Narathura japonica* (Murray) to learn how to recognize this mutualistic partner whereas they do not do the same with the non-ant-associated lycaenid, *Lycaena phlaeas* (Linnaeus) (Hojo *et al.*, 2014b). According to Hojo *et al.* (2014a) multiple chemical signatures, not only CHCs, may be important for a myrmecophilous caterpillar to exploit ants. Our study found high SIs among lycaenid species in unrelated genera. This convergence neither provides chemical camouflage on host plants nor chemical mimicry with tending ants. Thus, we propose ‘chemical conspicuousness’ as a strategy mediated by CHCs in myrmecophilous caterpillars. In this scenario, we can hypothesize the existence of Müllerian mimicry rings between myrmecophilous species with functional DNO — i.e. an honest signal between true mutualistic partners (Rossato & Kaminski, 2019). Batesian mimicry could also occur when caterpillars that invest little energy in secretions mimic CHCs of conspicuous caterpillars that secrete better caloric rewards for ants. A similar scenario proposed by Oliver & Stein (2011), was that caterpillar species lacking a DNO could chemically mimic rewarding caterpillars to gain protection by tending ants. Extensive comparative studies can reveal whether these chemical strategies do indeed occur at the community level.

The present study, along with others, reinforce the selective importance of chemical camouflage for herbivorous insects living on host plants frequently visited by ants (Akino, 2008; Silveira *et al.*, 2010). Chemical camouflage may have arisen as a defensive strategy allowing caterpillars to coexist with ants on plants. In multimodal systems with signals and cues between caterpillars and ants (see Casacci *et al.*, 2019), CHC composition can play a key adaptive role in decreasing ant attacks and increasing caterpillar protection and, consequently, survival. Myrmecophilous caterpillar assemblages can be quite rich in species that use different plant resources that vary in terms of nutritional quality and enemy-free space (Seufert & Fiedler, 1996; Rodrigues *et al.*, 2010; Silva *et al.*, 2014). Our study adds a new adaptive layer in the form of variation in the degree of similarity and camouflage efficiency that different host plants can offer myrmecophilous caterpillars. We hope that our study can serve as an incentive for further studies of the chemical interface of caterpillar-ant-plant interactions.

## Acknowledgments

We thank Laboratório Síncrotron for allowing us to work in its area of cerrado, and Instituto de Botânica de São Paulo for granting permission to work at Reserva Biológica and Estação Experimental de Mogi-Guaçu. Special thanks go to André V. L. Freitas and Paulo S. Oliveira for logistic support at UNICAMP and encouragement to study the chemical interface of plant-ant-butterfly interactions. Daniela Rodrigues and Adilson Moreira kindly assisted with the fieldwork. We are grateful to Sabrina C. Thiele for providing the photo of *Michaelus thordesa* and to Fábio S. do Nascimento, Geraldo L. G. Soares and Viviane G. Ferro for critically reviewing the manuscript. This study was financed by Coordenação de Aperfeiçoamento de Pessoal de Nível Superior – Brasil (CAPES) – Finance Code 001 as a PhD grant to LDL. JRT acknowledges grants from Fundação de Amparo à Pesquisa do Estado de São Paulo (FAPESP) (2011/17708-0) and Conselho Nacional de Pesquisa (CNPq) (2009/304473-0). LAK acknowledges grants from CNPq (140183/2006-0), FAPESP (10/51340-8), PNPD-CAPES, and the National Geographic Society (#WW-224R-17).

## Contribution of authors

All authors contributed to the study conception and design. LDL carried out the statistical analyses and wrote the manuscript. JRT carried out the chemical analyses and drafted the study. LAK collected the organisms, coordinated the study and critically revised the manuscript. All authors gave final approval for publication.

## Data availability statement

The data that support the findings of this study are openly available at https://doi.org/10.6084/m9.figshare.12584765.

## Supporting Information

**Table S1.**
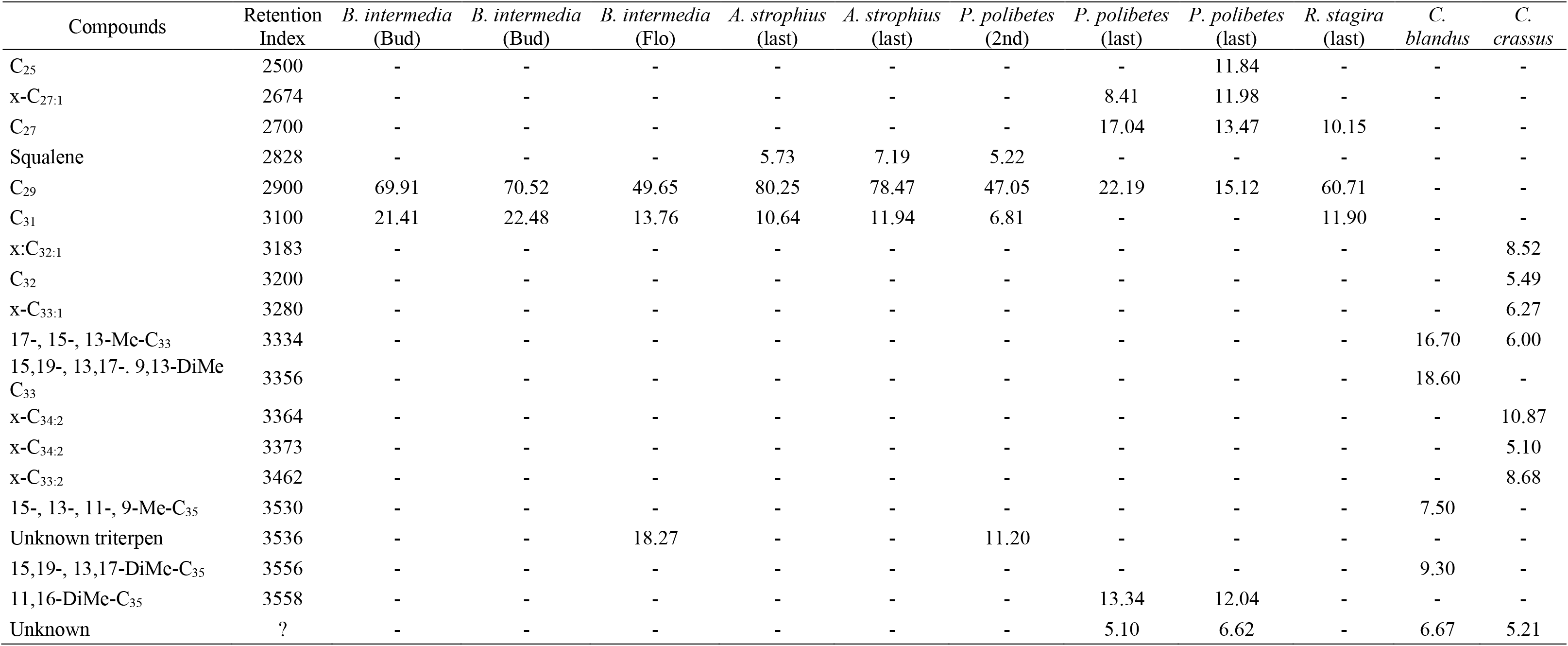
Relative abundance (%) of cuticular compounds of the host plant *Byrsonima intermedia*, lycaenid caterpillars and ants. Only compounds and values above 5% are shown

| Compounds | Retention Index | *B. intermedia* (Bud) | *B. intermedia* (Bud) | *B. intermedia* (Flo) | *A. strophius* (last) | *A. strophius* (last) | *P. polibetes* (2nd) | *P. polibetes* (last) | *P. polibetes* (last) | *R. stagira* (last) | *C. blandus* | *C. crassus* |
| --- | --- | --- | --- | --- | --- | --- | --- | --- | --- | --- | --- | --- |
| C<sub>25</sub> | 2500 | - | - | - | - | - | - | - | 11.84 | - | - | - |
| x-C<sub>27:1</sub> | 2674 | - | - | - | - | - | - | 8.41 | 11.98 | - | - | - |
| C<sub>27</sub> | 2700 | - | - | - | - | - | - | 17.04 | 13.47 | 10.15 | - | - |
| Squalene | 2828 | - | - | - | 5.73 | 7.19 | 5.22 | - | - | - | - | - |
| C<sub>29</sub> | 2900 | 69.91 | 70.52 | 49.65 | 80.25 | 78.47 | 47.05 | 22.19 | 15.12 | 60.71 | - | - |
| C<sub>31</sub> | 3100 | 21.41 | 22.48 | 13.76 | 10.64 | 11.94 | 6.81 | - | - | 11.90 | - | - |
| x:C<sub>32:1</sub> | 3183 | - | - | - | - | - | - | - | - | - | - | 8.52 |
| C<sub>32</sub> | 3200 | - | - | - | - | - | - | - | - | - | - | 5.49 |
| x-C<sub>33:1</sub> | 3280 | - | - | - | - | - | - | - | - | - | - | 6.27 |
| 17-, 15-, 13-Me-C<sub>33</sub> | 3334 | - | - | - | - | - | - | - | - | - | 16.70 | 6.00 |
| 15,19-, 13,17-. 9,13-DiMe C<sub>33</sub> | 3356 | - | - | - | - | - | - | - | - | - | 18.60 | - |
| x-C<sub>34:2</sub> | 3364 | - | - | - | - | - | - | - | - | - | - | 10.87 |
| x-C<sub>34:2</sub> | 3373 | - | - | - | - | - | - | - | - | - | - | 5.10 |
| x-C<sub>33:2</sub> | 3462 | - | - | - | - | - | - | - | - | - | - | 8.68 |
| 15-, 13-, 11-, 9-Me-C<sub>35</sub> | 3530 | - | - | - | - | - | - | - | - | - | 7.50 | - |
| Unknown triterpen | 3536 | - | - | 18.27 | - | - | 11.20 | - | - | - | - | - |
| 15,19-, 13,17-DiMe-C<sub>35</sub> | 3556 | - | - | - | - | - | - | - | - | - | 9.30 | - |
| 11,16-DiMe-C<sub>35</sub> | 3558 | - | - | - | - | - | - | 13.34 | 12.04 | - | - | - |
| Unknown | ? | - | - | - | - | - | - | 5.10 | 6.62 | - | 6.67 | 5.21 |

**Table S2.** Relative abundance (%) of cuticular compounds of the host plant *Pyrostegia venusta* and lycaenid caterpillars. Only compounds and values above 5% are shown

| Compounds | Retention Index | *P.venusta* (Bud) | *P. venusta* (Flo) | *M. thordesa* (last) | *P. polibetes* (last) | *P. polibetes* (last) | *P. polibetes* (last) | *P. polibetes* (last) | *R. marius* (last) |
| --- | --- | --- | --- | --- | --- | --- | --- | --- | --- |
| C<sub>23</sub> | 2300 | - | - | - | 12.84 | - | 5.56 | 9.58 | 12.00 |
| C<sub>25</sub> | 2500 | - | - | 25.66 | 18.39 | 23.53 | 19.55 | 11.17 | 7.06 |
| C<sub>26</sub> | 2600 | - | - | 6.07 | - | - | - | - | - |
| C<sub>27</sub> | 2700 | 15.02 | 20.18 | 36.09 | 28.69 | 27.76 | 29.67 | 23.72 | 12.71 |
| C<sub>28</sub> | 2800 | - | - | 6.52 | - | - | - | - | - |
| Squalene | 2828 | - | - | - | - | 7.72 | - | 12.50 | - |
| C<sub>29</sub> | 2900 | 33.28 | 48.37 | 13.03 | 12.22 | 13.76 | 17.50 | 14.75 | 20.09 |
| C<sub>31</sub> | 3100 | 23.37 | 16.81 | - | - | - | - | - | 20.66 |
| C<sub>33</sub> | 3300 | 5.16 | - | - | - | - | - | - | - |
| 15,19-, 13,17-DiMe-C<sub>35</sub> | 3556 | - | - | - | 6.70 | - | - | - | - |

**Table S3.**
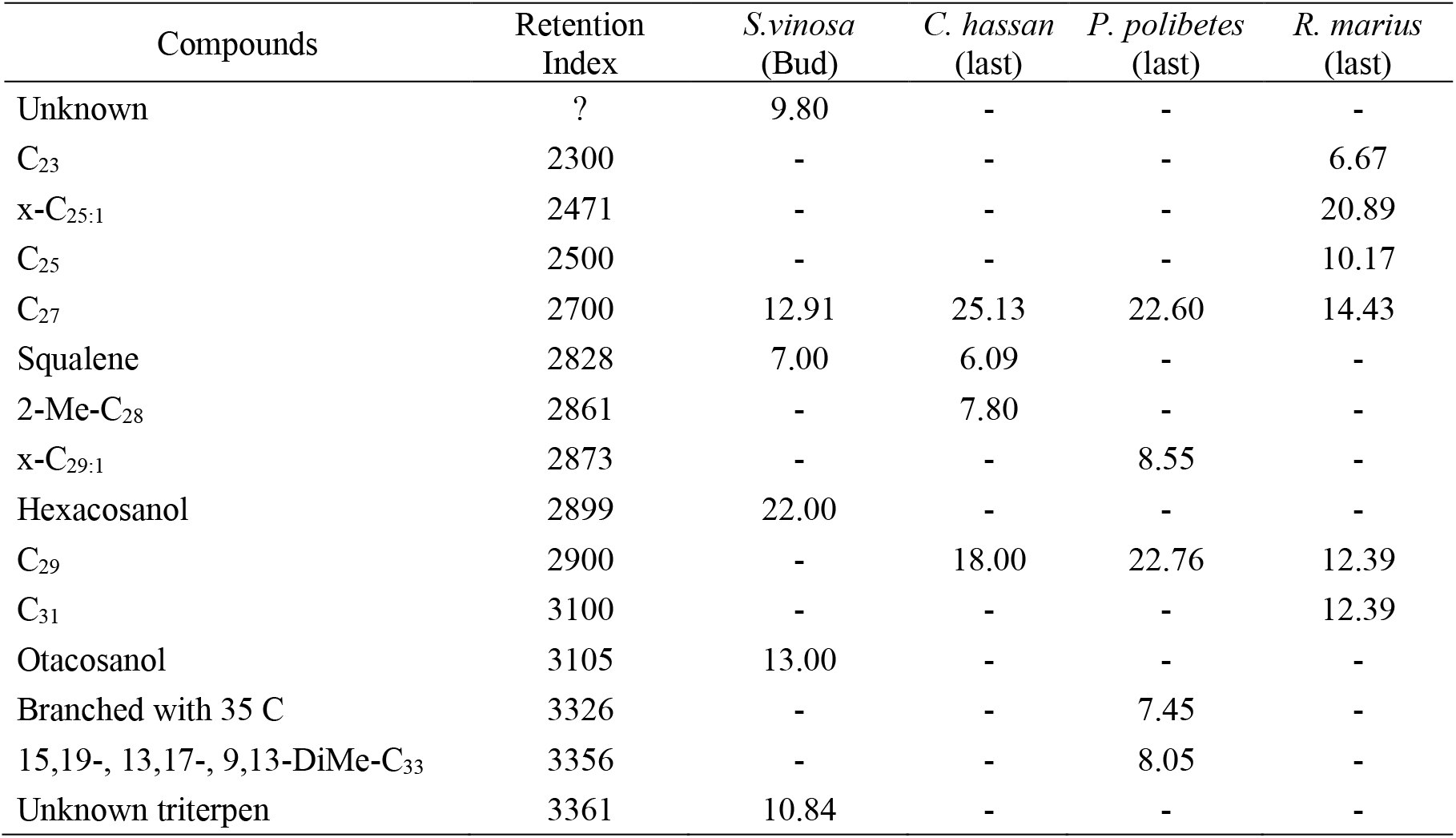
Relative abundance (%) of cuticular compounds of the host plant *Schefflera vinosa* and lycaenid caterpillars. Only compounds and values above 5% are shown

| Compounds | Retention Index | *S. vinosa*<br>(Bud) | *C. hassan*<br>(last) | *P. polibetes*<br>(last) | *R. marius*<br>(last) |
| --- | --- | --- | --- | --- | --- |
| Unknown | ? | 9.80 | - | - | - |
| C<sub>23</sub> | 2300 | - | - | - | 6.67 |
| x-C<sub>25:1</sub> | 2471 | - | - | - | 20.89 |
| C<sub>25</sub> | 2500 | - | - | - | 10.17 |
| C<sub>27</sub> | 2700 | 12.91 | 25.13 | 22.60 | 14.43 |
| Squalene | 2828 | 7.00 | 6.09 | - | - |
| 2-Me-C<sub>28</sub> | 2861 | - | 7.80 | - | - |
| x-C<sub>29:1</sub> | 2873 | - | - | 8.55 | - |
| Hexacosanol | 2899 | 22.00 | - | - | - |
| C<sub>29</sub> | 2900 | - | 18.00 | 22.76 | 12.39 |
| C<sub>31</sub> | 3100 | - | - | - | 12.39 |
| Otacosanol | 3105 | 13.00 | - | - | - |
| Branched with 35 C | 3326 | - | - | 7.45 | - |
| 15,19-, 13,17-, 9,13-DiMe-C<sub>33</sub> | 3356 | - | - | 8.05 | - |
| Unknown triterpen | 3361 | 10.84 | - | - | - |

**Table S4.** One-way ANOSIM comparing the relative proportions of cuticular hydrocarbons of the profiles of lycaenid caterpillars, host plants and workers of the genus *Camponotus*

|  | Caterpillars |  | Ants |  | Plants |  |
| --- | --- | --- | --- | --- | --- | --- |
|  | R | P | R | P | R | P |
| Caterpillars | - | - | 1 | 0.0067 | 0.2094 | 0.0329 |
| Ants | 1 | 0.0067 | - | - | 0.6354 | 0.0768 |
| Plants | 0.2094 | 0.0329 | 0.6354 | 0.0768 | - | - |

**Table S5.** One-way ANOSIM comparison between the relative proportions of cuticular hydrocarbons of the profiles of lycaenid caterpillars, host plants and *Camponotus* workers

|  | Commensal caterpillars |  | Mutualistic caterpillars |  | Ants |  | Plants |  |
| --- | --- | --- | --- | --- | --- | --- | --- | --- |
|  | R | P | R | P | R | P | R | P |
| Commensal caterpillars | - | - | 0.744 | 0.0048 | 1 | 0.0924 | -0.08642 | 0.5548 |
| Mutualistic caterpillars | 0.744 | 0.0048 | - | - | 1 | 0.0101 | 0.5535 | 0.0007 |
| Ants | 1 | 0.0924 | 1 | 0.0101 | - | - | 0.6354 | 0.068 |
| Plants | -0.08642 | 0.5548 | 0.5535 | 0.0007 | 0.6354 | 0.068 | - | - |

